# Time-dependent memory and individual variation in Arctic brown bears (*Ursus arctos*)

**DOI:** 10.1101/2021.12.09.472008

**Authors:** Peter R. Thompson, Mark A. Lewis, Mark A. Edwards, Andrew E. Derocher

## Abstract

Animal movement modelling provides unique insight about how animals perceive their landscape and how this perception may influence space use. When coupled with data describing an animal’s environment, ecologists can fit statistical models to location data to describe how spatial memory informs movement. We performed such an analysis on a population of brown bears (*Ursus arctos*) in the Canadian Arctic using a model incorporating time-dependent spatial memory patterns. Brown bear populations in the Arctic lie on the periphery of the species’ range, and as a result endure harsh environmental conditions. In this kind of environment, effective use of memory to inform movement strategies could spell the difference between survival and mortality. The model we fit tests four alternate hypotheses (some incorporating memory; some not) against each other, and we found a high degree of individual variation in how brown bears used memory. We found that 52% (11 of 21) of the bears used complex, time-dependent spatial memory to inform their movement decisions. These results, coupled with existing knowledge on individual variation in the population, highlight the diversity of foraging strategies for Arctic brown bears while also displaying the inference that can be drawn from this innovative movement model.

## 2 Introduction

Ecologists have used animal movement data to answer many important ecological questions in recent years (Nathan et al., 2008; Joo et al., 2020). Models have been developed to explore the qualities of an animal’s home range (Worton, 1989; Dahle and Swenson, 2003; Borger et al., 2008; Edwards et al., 2009), large-scale movements such as migration (Dingle and Drake, 2007; Merkle et al., 2019), and species-habitat relationships (i.e., habitat selection; Boyce and McDonald, 1999; Fortin et al., 2005; Thurfjell et al., 2014). Habitat selection analyses, in particular, have advanced due to the increasing availability of remote sensing data, which can describe large-scale environmental patterns, as well as animal movement data itself (Kays et al., 2015). These analyses provide solutions to difficult problems concerning how animals interact with their environment (Muhly et al., 2019; Suraci et al., 2019). Understanding these interactions, however, is limited without incorporating how animals perceive their environments cognitively (Fagan et al., 2017). This realization in movement ecology has inspired the growth of memory-informed movement modelling.

By including spatial memory, we can quantitatively model animal cognition using movement data. Animals use spatial memory to encode, store, and retrieve information about the location of landmarks in an animal’s environment (Fagan et al., 2013). Ecologists have included memory into habitat selection models by hypothesizing that animals will select for areas they have visited more frequently (Dalziel et al., 2008; Oliveira-Santos et al., 2016), assuming animals will select against areas they have just visited (Schlägel and Lewis, 2014), or modifying habitat selection models such that animals will not be attracted to high-quality patches unless they can perceive this quality (Van Moorter et al., 2009; Avgar et al., 2013). Most of these models lack attention to temporal memory, where animals remember not just where they have visited but how long ago they were there. While the “time since last visit” construct incorporated by Schlägel and Lewis (2014) is a noteworthy exception, they assumed patches become increasingly attractive to the animal as time passes, which is not realistic in seasonally variable environments. For animals with seasonally varying home ranges, the energetic value of visiting a food patch may vary periodically or seasonally. Animals that live in such environments may change the size and shape of their home range seasonally, implying that they only visit specific parts of their home range at specific times of year (Wiktander et al., 2001). On a smaller timescale, spatiotemporal memory allows animals to capitalize on ripe fruit, which loses its energetic return if visited too late (Janmaat et al., 2016). Despite the occurrence of such patterns, which may be either ephemeral or seasonal, animal movement models rarely incorporate a time-dependent spatial memory mechanism that accounts for them.

The brown bear (*Ursus arctos*) is a widespread, omnivorous mammal found in the Northern Hemisphere (Pasitschniak-Arts, 1993), and populations in seasonal regions of the species’ range are likely to benefit from remembering the timing of food resources. The Canadian Arctic is an example of such an environment, and brown bears that live here are especially opportunistic, taking advantage of a wide variety of food resources (Edwards and Derocher, 2015). Most brown bear food resources here are only available for a fraction of the bears’ active season (Nagy and Haroldson, 1990; Burn and Kokelj, 2009; Edwards and Derocher, 2015), resulting in seasonal variation in their habitat selection (McLoughlin et al., 2002). Brown bears in the Arctic also display individual dietary variation due to sexual size dimorphism as well as the reproductive constraints of adult females (Edwards et al., 2011). Theoretical studies have displayed the utility of memory-informed movement in environments with predictable temporal variation (Mueller et al., 2011). Evidence of memory-informed movement in other brown bear populations includes oriented movement towards previously visited kill sites (Selva et al., 2017), scent marking to identify territorial boundaries (Clapham et al., 2012), fidelity to the same salmon-rich stream each year (Wirsing et al., 2018), and repeated use of the same denning area each year (Manchi and Swenson, 2005; Sorum et al., 2019). These studies demonstrate the cognitive and perceptual capabilities of the species, suggesting that brown bears in the Canadian Arctic may incorporate time-dependent spatial memory into their movement patterns.

We applied a new animal movement model that incorporates a unique form of complex, time-dependent spatial memory (Thompson et al., 2021) to global positioning system (GPS) data for brown bears from the Mackenzie Delta region of the Canadian Arctic. Thompson et al. (2021) designed a model with four special cases, each concerning its own hypothesis about cognition and movement: a null hypothesis; a resource-only hypothesis assuming simple resource selection; a memory-only hypothesis assuming resource-less seasonal revisitation patterns within an animal’s home range; and a resource-memory hypothesis assuming animals are simultaneously influenced by local resources and spatial memory. Fitting each of these four models to animal location data provides inference on the relative likelihood of each hypothesis being true, and the parameters in each model describe explicit components of the animal’s foraging behaviour. We obtained parameter estimates and performed model selection analysis for 21 individual bears, allowing us to explicitly examine variation at the individual level. We found that amid high individual variation within the population, movement patterns from a majority of the bears supported the resource-memory hypothesis. These results represent the first application of a novel model to a population of opportunistic and potentially sensitive omnivores.

## 3 Materials and Methods

We applied the model described in Thompson et al. (2021) to global positioning system (GPS) location data from a population of brown bears in the Canadian Arctic. We used the model to test four alternate hypotheses stated above about animal movement and cognition (Figure 1). We drew inference from maximum likelihood estimates for the model parameters to quantify characteristics of the bears’ behaviour (Table 1). We describe the biological function of the model here, noting that it is described in full detail in Thompson et al. (2021).

**Figure 1:**
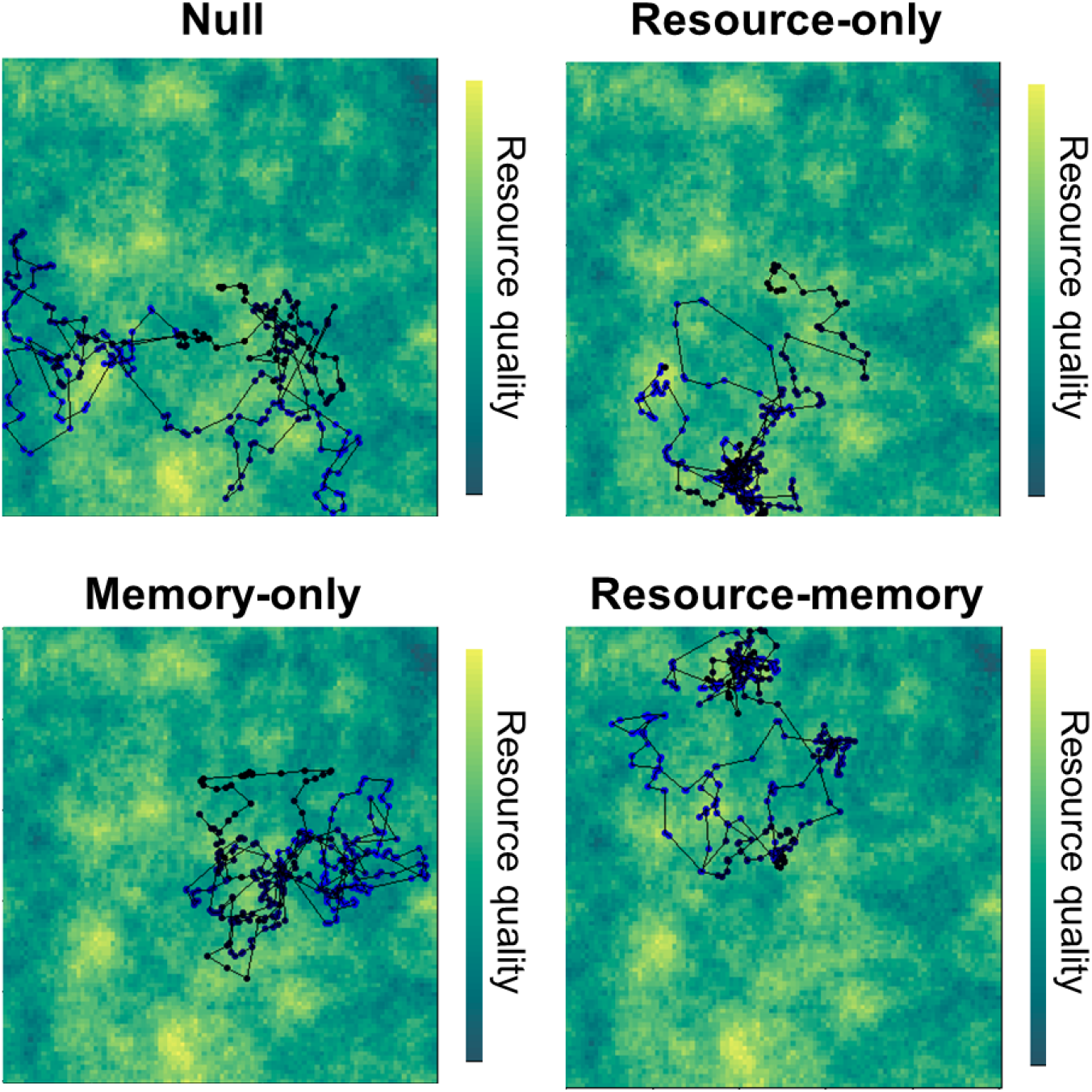
Simulated animal movement tracks (300 steps per track) on a randomly generated landscape displaying behaviours consistent with each hypothesis (and model). The colour of each point on this simulated movement track represents the hypothetical time in the animal’s memory “cycle”, which is here set to 100 time units (points at *t* = 75 have the same colour as *t* = 175). The null model implies completely random movement, while the resource-only model implies that the animal will locate nearby resources and select for those areas. The memory-only model implies that the animal relocates itself to areas it visited 100 time units before. The resource-memory model combines mechanisms in the resource-only and memory-only models.

**Table 1:** Description of model parameters, including units (N/A implies that the parameter is unitless) and models (N = null; R = resource-only; M = memory-only; RM = resource-memory) in which the parameters are estimated. Adapted from Thompson et al. (2021).

|  | Units | Description | N | R | M | RM |
| --- | --- | --- | --- | --- | --- | --- |
| $\rho_{ns}$ | km/h | Mean movement speed in non-stationary state | X | X | X | X |
| $\kappa$ | N/A | Degree of directional autocorrelation | X | X | X | X |
| $\beta_0$ | N/A | Probability of revisitation | | | | X |
| $\beta_1$ | N/A | Selection coefficient for berries | | X | | X |
| $\beta_2$ | $\frac{1}{km}$ | Selection coefficient for riparian habitats | | X | | X |
| $\beta_3$ | N/A | Selection coefficient for squirrels | | X | | X |
| $\beta_4$ | N/A | Selection coefficient for sweetvetch | | X | | X |
| $\beta_5$ | $\frac{1}{km}$ | Selection coefficient for human settlements | | X | | X |
| $\beta_6$ | $\frac{1}{km}$ | Selection coefficient for cabins | | X | | X |
| $\beta_d$ | N/A | Strength of selection for memorized areas | | | X | X |
| $\mu$ | days | Mean time lag between revisitations | | | X | X |
| $\sigma$ | days | Standard deviation in time between revisitations | | | X | X |
| $\alpha$ | $\log(km)$ | Degree of perceptual resolution | | | X | X |
| $\lambda$ | N/A | Probability of staying in stationary state | X | X | X | X |
| $\gamma$ | N/A | Probability of staying in non-stationary state | X | X | X | X |

### 3.1 Study area

The Mackenzie River empties into the Arctic Ocean in the northern Northwest Territories, in NW Canada. Our study area, the Mackenzie Delta region, spans 23,000 km^2^ of wet Arctic tundra, interspersed with many lakes and smaller streams (Edwards et al., 2013). The Mackenzie Delta region is a harsh environment for brown bears, with minimal food availability that results in short active seasons (Ferguson and McLoughlin, 2000). There are two human settlements in the region, Inuvik and Tuktoyaktuk, in addition to some remote and rarely inhabited industrial camps.

Our landscape data provide information on the spatial heterogeneity in vegetation and topography. We used three 30 × 30 m raster layers to describe the study area: a digital elevation model (DEM) measuring elevation (ranging from 0 m to 1676 m), a vegetation class raster describing dominant vegetation in each portion of the landscape, and a raster approximating the density of Arctic ground squirrels (*Urocitellus parryii*), which are a common brown bear prey species (MacHutchon and Wellwood, 2003; Barker et al., 2015). The vegetation class raster classified each 30 × 30 m grid cell into one of 46 vegetation classes, describing the age, size, and/or dominant plant species present in each area (Ducks Unlimited, 2002; but also see the Appendix). The ground squirrel raster is a product of a resource selection function from an existing study, so it quantifies the likelihood (based on environmental conditions) for any spatial region to support ground squirrels (Barker and Derocher, 2010).

### 3.2 Brown bear data

Between 2003, and 2006, 31 brown bears (24 female, 7 male) were captured and equipped with GPS collars (Telconics Inc., Mesa, AZ, USA) that provided the bear’s spatial location every four hours. The collars used long temporal sequences without movement to identify denning periods, and did not record any signals until the bear began to move again in the spring. The collars were removed and/or stopped recording bear locations after one to four years. The University of Alberta Animal Care and Use Committee for Biosciences approved all animal capture and handling procedures, which were in accordance with the Canadian Council on Animal Care. Capture was conducted under permit from the Government of the Northwest Territories.

### 3.3 Model design

We fit a discrete-time hidden Markov model (HMM) that assesses the nature of complex time-dependent spatial memory mechanisms in Arctic brown bears. The model has two movement states: one representing resting or not moving (stationary), and one representing movement (non-stationary). In a HMM, the state is not explicitly known but can be inferred from observed data (e.g., if consecutive GPS locations are only 1 m apart, we can infer that the bear is likely in the stationary state), which is mathematically expressed with conditional likelihood functions (Jonsen et al., 2013). Like other HMMs, the bear’s movement state at any point in time depends only on the previous state as well as fixed state-switching probabilities. For a two-state HMM, only two probabilities are necessary to explain the entire system: λ, the probability of remaining in the stationary state, and *γ*, the probability of remaining in the non-stationary state. We fit both of these values as model parameters (Thompson et al., 2021).

When the bear is in the non-stationary state, the probability distribution of its steps, which we denote *f_ns_*, strongly resembles a step selection function (Fortin et al., 2005). Each step consists of two consecutive locations (e.g., **x**_*t*–1_ to **x**_t_), but we must also consider the heading on which the animal arrived at **X**_*t*–1_, *ϕ*_*t*–1_. It is a normalized product of two components: *k*, the resource-independent movement kernel, and *W*, the environmental (or cognitive) weighting function:

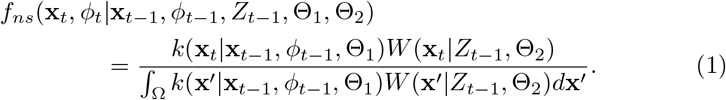

Here *k* depends on the animal’s previous location, its previous heading, and Θ_1_, a set of model parameters corresponding to resource-independent movement. The functional form of *k*, which resembles a correlated random walk, is the same in each model (Thompson et al., 2021). The parameter vector Θ_2_ contains all parameters concerning cognition or habitat selection, and because Θ_2_ contains different parameters in each model, we introduce Θ_2,*N*_, Θ_2,*R*_, Θ_2,*M*_, and Θ_2,*RM*_ for the null, resource-only, memory-only, and resource-memory models, respectively (we also introduce *W_N_*, *W_R_*, *W_M_*, and *W_RM_* for the same reasons). The weighting function *W* also depends on *Z*_*t*–1_, the animal’s cognitive map at time *t* – 1, which quantifies and spatially organizes the animal’s past experiences and memories.

In practice, we approximated the denominator of Equation 1 with a sum, as is standard with step selection functions (Fortin et al., 2005; Thurfjell et al., 2014), by simulating a set of “available steps” from *k* and calculating *W* at those steps (Thompson et al., 2021). The most accurate combination of model parameters is the one such that the animal’s observed steps register significantly higher values of *W* than would be expected from random movement (i.e., *k*).

#### 3.3.1 Null model

In the null model, we assume that the bear moves randomly, so there are only four parameters of concern: λ, *γ, ρ_ns_*, and *κ* (Table 1). If the 95% confidence interval for *κ* excludes 0, we can conclude that there exists significant directional autocorrelation in the bear’s movements. The weighting function *W_N_*(*x_t_*|*Z*_*t*–1_, Θ_2,*N*_) = 1 for all *x_t_* in space (note that Θ_2_ is just an empty vector here), making *f_ns_* equal to *k* in the null model.

#### 3.3.2 Resource-only model

The resource-only model tests the hypothesis that bears select for nearby locations with high habitat quality. *W_R_* resembles a step selection function and Θ_2,*R*_ contains a vector of selection coefficients (*β*_1_,*β*_2_, …, *β_P_*) for each resource covariate (*r*_1_(*x*), *r*_2_(*x*), …, *r_P_*(*x*)) in the model. We define *W_R_* as follows:

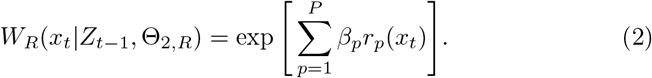

If the 95% confidence interval for any of these parameters excludes 0, we can conclude that the animal significantly selected for (or against) that variable.

We manipulated our landscape data to produce *P* = 6 resource covariates. Berries (including but not limited to *Empetrum nigrum, Shepherdia canadensis, Vaccinium uliginosum*, and *V. vitis-idaea*) are an important dietary item for most individuals (Edwards et al., 2011; Edwards and Derocher, 2015). In the Canadian Arctic, berries are generally found in dwarf shrub areas (Porsild and Cody, 1980; Shevtsova et al., 1995; Norment and Fuller, 1997), but they can also occur beneath the canopy of northern woodlands (Murray et al., 2005). We do not have an explicit berry density survey, so we used the vegetation class data along with knowledge of common berry species to infer the probability of berries occurring at each spatial grid cell (Table **??**).

We included a covariate representing the Euclidean distance from turbid water to gauge the extent to which brown bears select for riparian areas. These regions support food resources such as horsetails (*Equisetum* spp.) and wetland sedges (*Carex* spp.) that are important in the early summer (Edwards and Derocher, 2015). Brown bears in the Mackenzie Delta region also fish for broad whitefish (*Coregonus nasus*) beside streams and rivers when the fish migrate (Barker and Derocher, 2009).

We also included covariates representing the possible presence of Arctic ground squirrels and alpine sweetvetch (*Hedysarum alpinum*), two common dietary resources in the area (Edwards and Derocher, 2015). We used the ground squirrel RSF from Barker and Derocher (2010) as a covariate for squirrel selection. Sweetvetch occurs in dry, shrubby uplands (Porsild and Cody, 1980), so we used an interaction between slope (from our DEM) and dwarf shrub vegetation classes to quantify sweetvetch density.

Brown bears are affected by the presence of humans in many ways (Mace et al., 1996; Steyaert et al., 2016; Lamb et al., 2017), so we included covariates measuring the Euclidean distance from various human settlements or dwellings. The first covariate measured the distance from the nearest human settlement in the Mackenzie Delta region (either Inuvik or Tuktoyaktuk). Brown bears that come near human settlements are often deterred by the residents or wildlife officials in a forceful manner (Kellert et al., 1996), so we expected bears whose home ranges overlap one of the settlements to avoid them. Some more remote industrial buildings are occasionally inhabited but often lack the constant human presence brown bears face near Inuvik or Tuktoyaktuk. As opportunistic omnivores, brown bears commonly use anthropogenic food sources (Kavčič et al., 2015) and may visit these buildings. Our second anthropogenic covariate measured the Euclidean distance from the closest of the 6 cabins in the region.

#### 3.3.3 Memory-only model

The memory-only model quantifies the hypothesis that brown bears remember the spatial location of areas they have visited previously, with the intent to return there after a consistently scheduled time lag (Thompson et al., 2021). The cognitive map associated with this model builds on the idea of time since last visit proposed by Schlägel and Lewis (2014), where previously visited locations become increasingly more attractive to the animal as time increases. We model this structure with a discrete-space cognitive map *Z_t_* where the animal keeps track of all its previous locations. For any spatial grid cell *z* on the map, *Z_t_*(*z*) contains a linked list of time since previous visits to *z*. This structure allows the memory of multiple visits to the same location.

The memory-only model follows the hypothesis that there is some “peak” in attractiveness that represents the periodicity of habitat quality in the environment. We fit the timing of this peak *μ*, as well as the degree of concentration and variation around this peak *σ*, as model parameters. Higher values of *σ* indicate that bears are less precise in their revisitation patterns, and may also be indicative of lower temporal predictability in the environment. Specifically, *W_M_* is a weighted average of distances from previously visited locations on the bear’s track, where the weights correspond to Gaussian distribution values with parameters *μ* and *σ*. For each time lag *τ* we can use *Z_t_* to identify where the bear was at time *t* – *τ* (let us denote this location *z_t–τ_*). Then, the weight for each point is equal to *φ*(*τ*|*μ*, *σ*), where *φ* is the Gaussian probability distribution function.

Each distance is transformed exponentially with parameter *α*. For smaller values of *α*, the mathematical difference between a step 1 km away and a step 2 km away would be amplified, suggesting that the bear would interpret these spatial differences more dramatically. If the 95% confidence interval for *α* is entirely below 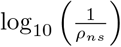, then the bear perceives its landscape as relatively heterogeneous (the opposite is true if the confidence interval is entirely above this value). The memory-only model includes one last parameter *β_d_*, representing the relative probability of revisiting a memorized location instead of moving randomly or selecting for present-time resources. As *β_d_* approaches 1, the animal will approach oriented movement towards previously visited locations, and if the 95% confidence interval for this parameter excludes 0.5, we can conclude that the animal is displaying significant selection for memorized areas.

We formulate *W_M_* as follows:

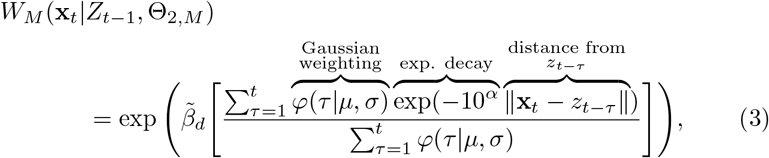

where 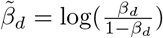.

#### 3.3.4 Resource-memory model

The resource-memory model combines the principles of the resource-only and memory-only models (Thompson et al., 2021). Bears moving according to this model consider present-time resources in nearby locations as well as previously visited locations. We additionally hypothesize that bears will only be attracted to previously visited locations that had food, and will not revisit previously visited locations with low resource quality. This mechanism is mediated by “threshold” parameter *β*_0_, which represents the relative probability of returning to previously visited locations. If *β*_0_ = 1 then the animal perceives all previously visited locations as “attractive” for revisitation, regardless of habitat quality, and if *β*_0_ = 0 then the opposite is true. We can infer about the habitat quality necessary to influence revisitations from a bear if the 95% confidence interval for *β*_0_ overlaps 0.5, which would imply being neutrally selective towards these locations.

The weighting function for the resource-memory model includes two terms, one representing present-time resource selection and one representing memorized information:

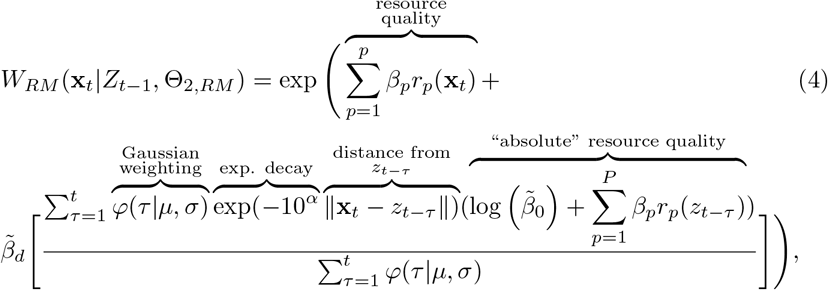

where 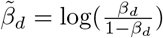 and 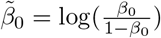.

### 3.4 Fitting the model to data

We used maximum likelihood estimation to fit the four models to each individual, comparing each model using the Bayesian Information Criterion (BIC). Thompson et al. (2021) found that BIC was more accurate than AIC in terms of selecting the most parsimonious model for data. BIC allowed us to identify the “best model”, or the model associated with the hypothesis that is most likely to be true, for each bear. A difference in information criteria greater than 2 between the best and second-best models indicates greater support for the best model (Burnham and Anderson, 2004). We used maximum likelihood estimates (MLEs) along with 95% confidence intervals for each parameter in the best model to obtain further information on the bears’ movement behaviours. We removed the first year of GPS data from model fitting for every bear because we did not have the necessary information of previous locations for that portion of the data. We refer to this first year as “training data”, and removed bears with only one year of GPS data from the analysis. The models are computationally complex, so we used advanced automatic differentiation techniques to obtain MLEs (Albertsen et al., 2015; Kristensen et al., 2016; Whoriskey et al., 2017) and likelihood profiling to obtain confidence intervals (Fischer and Lewis, 2020). See Thompson et al. (2021) as well as the Appendix for additional details on model fitting.

#### 3.4.1 Seasonal resource modelling

To ensure that the seasonal revisitation patterns we observed were a result of spatial memory, we tested an alternate version of the model where resource covariates were restricted to seasons of availability. In the original versions of the resource-only and resource-memory model, each resource covariate *r_i_*(*x*) retains the same value throughout the year. This follows the assumption that our covariates measure the habitat conditions necessary to support seasonally available resources, not the resources themselves. For example, *r*_2_(*x*), the distance from *x* to the nearest riparian area, does not change seasonally, but the likelihood of obtaining valuable food resources from that region does vary seasonally. That being said, identifying memory based solely on movement patterns requires rigorously eliminating any other mechanisms that could cause those patterns (Fagan et al., 2013), so we designed an alternate model where resources were explicitly seasonal.

We identified temporal intervals in which each resource would be treated as present on the landscape, and assumed that *r_i_*(*x*) would be equal to 0 outside of these intervals. Berries are available in smaller portions year-round (Edwards and Derocher, 2015), but the primary period of occurrence lasts from approximately August 1 until the end of the active season, which we considered to be November 30, when bears had entered dens and GPS collars turned off (Gau et al., 2002; MacHutchon and Wellwood, 2003). The food available in riparian habitats (including whitefish, which generally migrate in early October; Barker and Derocher, 2009) is most prominent from May 10 to October 16 (Macdonald et al., 1995) when the ice has melted from the Mackenzie River. Arctic ground squirrels are always present, but they are easier for brown bears to hunt when they are hibernating (Barker et al., 2015), so we used an interval from September 11 to November 30 to approximate when most squirrels would be dormant (Buck and Barnes, 1999). Sweetvetch is also available year-round, but provides the highest nutritional return in the early spring, so we used an interval from April 1 (the beginning of the active season) to June 15 (MacHutchon and Wellwood, 2003). We left *r*_5_ and *r*_6_, the covariates relating to presence of humans, temporally constant.

We fit the models again with these newly defined resource covariates, performing all the same model selection and parameter estimation analyses. The null and memory-only models were formulated the same in both versions of the model fitting process. We used a separate set of “available points” (used to approximate Equation 1) for these analyses, so the outputs (BIC and parameter estimates) for these models were still slightly different.

## 4 Results

Of the 31 bears for which we had GPS data, 21 (18 females, 3 males) bears had enough data (at least one year excluding the first year of training) for model fitting. We fit all four models to each bear and used BIC to identify which associated hypothesis was most heavily supported by the data. Once we identified the best model for each bear, we calculated MLEs and 95% confidence intervals for all parameters in that model. We found that despite a large degree of individual variation, bears generally exhibit movement informed by resources as well as memory, with a revisitation scale close to 365 days. We also confirmed that memory, not the seasonality of resources, was the primary mechanism causing brown bears to return to previously visited food patches in a periodic fashion.

### 4.1 Model selection

The Mackenzie Delta brown bear population displayed a variety of movement behaviours, although the resource-memory model was most frequently selected as the most parsimonious explanation of the bears’ movement patterns (Table 2). It was identified as the “best model” (using BIC) for 9 of the 21 bears. The resource-only and memory-only models also received some support within the population; these models were the best model for 5 and 4 bears, respectively. For 3 of 21 bears, the null model was the most parsimonious explanation of bear movement patterns. There were only two cases where the difference in BIC between the two best models was < 2 (Table 2).

**Table 2:** dBIC (difference in BIC from the “best model”) values for each model and bear, with resource covariates set to be temporally constant. Cells shaded gray represent the model that best explains the movement patterns of each bear (dBIC = 0), and cells shaded light gray represent models < 2 BIC above the best model. Bears are sorted in descending order by number of data points (i.e., bears with more data at the top of the table).

| Bear ID | Null | Resource-only | Memory-only | Resource-memory |
| --- | --- | --- | --- | --- |
| GF1004 | 38.5 | 32.5 | 50.8 | 0.0 |
| GM1046 | 119.6 | 40.3 | 73.5 | 0.0 |
| GF1008 | 62.5 | 17.5 | 74.0 | 0.0 |
| GF1086 | 90.4 | 0.0 | 88.2 | 49.3 |
| GF1016 | 15.3 | 6.0 | 0.0 | 21.6 |
| GF1041 | 125.5 | 45.7 | 107.3 | 0.0 |
| GF1007 | 222.6 | 0.0 | 221.1 | 33.6 |
| GF1130 | 0.0 | 12.9 | 17.4 | 24.5 |
| GF1005 | 30.7 | 0.0 | 8.4 | 5.2 |
| GF1096 | 50.3 | 0.0 | 3.9 | 46.6 |
| GF1167 | 39.1 | 28.5 | 46.0 | 0.0 |
| GF1079 | 119.3 | 12.2 | 108.8 | 0.0 |
| GF1089 | 10.5 | 0.0 | 3.4 | 1.9 |
| GF1141 | 25.2 | 43.2 | 0.0 | 46.6 |
| GF1133 | 54.3 | 60.7 | 61.0 | 0.0 |
| GF1087 | 32.8 | 44.4 | 0.0 | 13.6 |
| GF1108 | 0.0 | 21.0 | 11.3 | 24.3 |
| GF1143 | 4.7 | 6.7 | 0.0 | 24.1 |
| GF1147 | 90.6 | 14.5 | 65.6 | 0.0 |
| GF1092 | 30.2 | 3.6 | 28.2 | 0.0 |
| GF1146 | 0.0 | 14.9 | 0.4 | 3.5 |

#### 4.1.1 Seasonal resource modelling

When we revised our resource covariates by adding time dependence, the memory-only model was a much more parsimonious explanation of the data (Table 3). It was the “best model” for 14 of the 21 bears when resource covariates were restricted to our prescribed seasons. The resource-only model was the best model for three bears, and the null and resource-memory model were the best for two bears each. There were three cases where the difference in BIC between the best model and the other models was < 2 (Table 3).

**Table 3:**
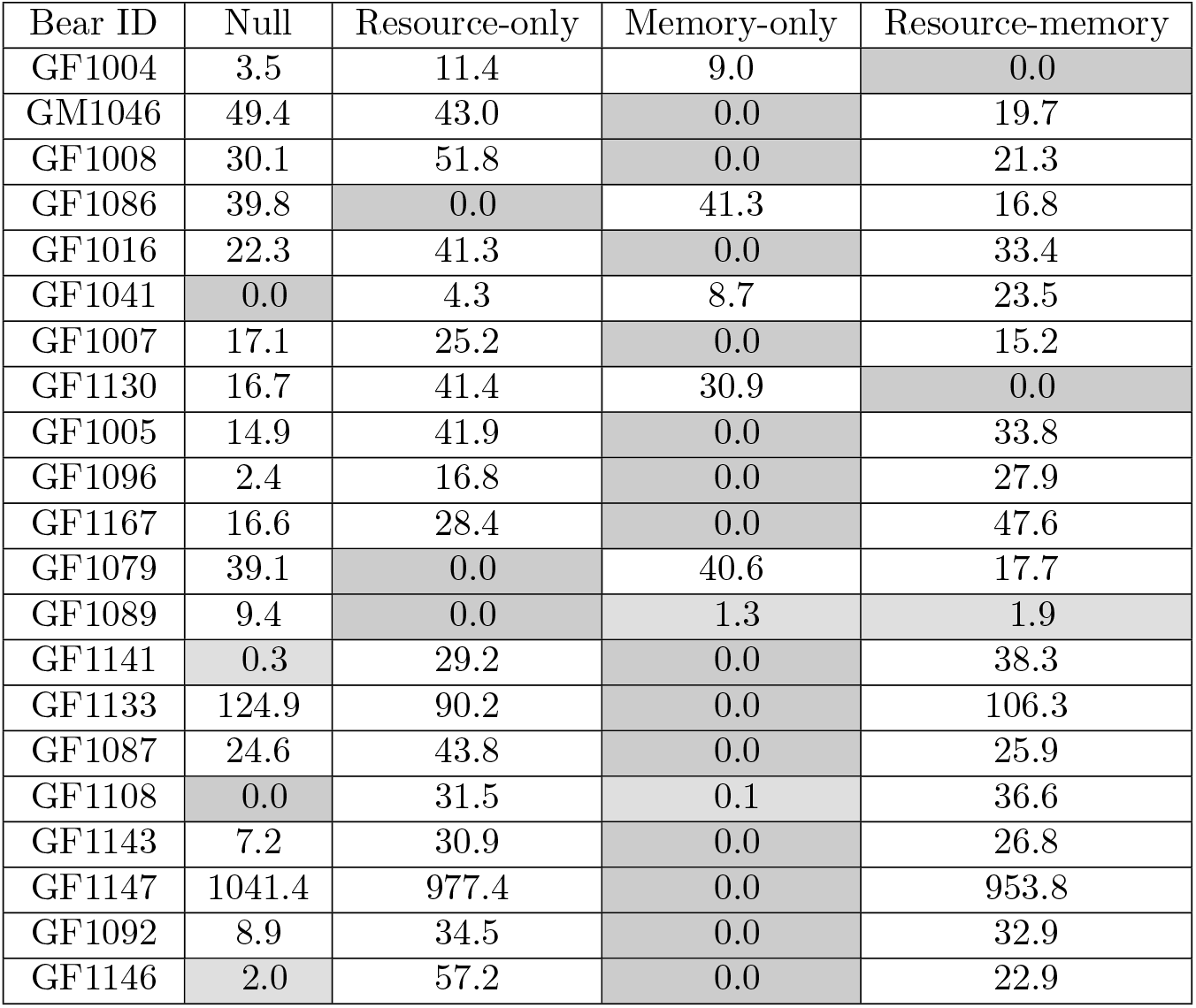
dBIC (difference in BIC from the “best model”) values for each model and bear, with resource covariates set to be temporally variable. Cells shaded gray represent the model that best explains the movement patterns of each bear (dBIC = 0), and cells shaded light gray represent models < 2 BIC above the best model. Bears are sorted in descending order by number of data points (i.e., bears with more data at the top of the table).

| Bear ID | Null | Resource-only | Memory-only | Resource-memory |
| --- | --- | --- | --- | --- |
| GF1004 | 3.5 | 11.4 | 9.0 | 0.0 |
| GM1046 | 49.4 | 43.0 | 0.0 | 19.7 |
| GF1008 | 30.1 | 51.8 | 0.0 | 21.3 |
| GF1086 | 39.8 | 0.0 | 41.3 | 16.8 |
| GF1016 | 22.3 | 41.3 | 0.0 | 33.4 |
| GF1041 | 0.0 | 4.3 | 8.7 | 23.5 |
| GF1007 | 17.1 | 25.2 | 0.0 | 15.2 |
| GF1130 | 16.7 | 41.4 | 30.9 | 0.0 |
| GF1005 | 14.9 | 41.9 | 0.0 | 33.8 |
| GF1096 | 2.4 | 16.8 | 0.0 | 27.9 |
| GF1167 | 16.6 | 28.4 | 0.0 | 47.6 |
| GF1079 | 39.1 | 0.0 | 40.6 | 17.7 |
| GF1089 | 9.4 | 0.0 | 1.3 | 1.9 |
| GF1141 | 0.3 | 29.2 | 0.0 | 38.3 |
| GF1133 | 124.9 | 90.2 | 0.0 | 106.3 |
| GF1087 | 24.6 | 43.8 | 0.0 | 25.9 |
| GF1108 | 0.0 | 31.5 | 0.1 | 36.6 |
| GF1143 | 7.2 | 30.9 | 0.0 | 26.8 |
| GF1147 | 1041.4 | 977.4 | 0.0 | 953.8 |
| GF1092 | 8.9 | 34.5 | 0.0 | 32.9 |
| GF1146 | 2.0 | 57.2 | 0.0 | 22.9 |

### 4.2 Parameter estimation

Most of the results below concern the “traditional” model, where we did not explicitly set the seasonality for the resource parameters. See Table **??** in the Appendix for parameter estimates when resources were explicitly seasonal.

#### 4.2.1 Movement parameters

Brown bears varied in their movement speed and directional autocorrelation (Table 4). Mean movement speed in the non-stationary state (*ρ_ns_*) varied from 0.22 (GF1086) to 0.65 (GF1005) km/h. Parameter estimates for *κ* varied from 0 (GF1092) to 0.7385 (GF1143), and 18 of the 21 bears exhibited some significant directional autocorrelation (i.e., the 95% confidence interval for *κ* excluded 0).

**Table 4:**
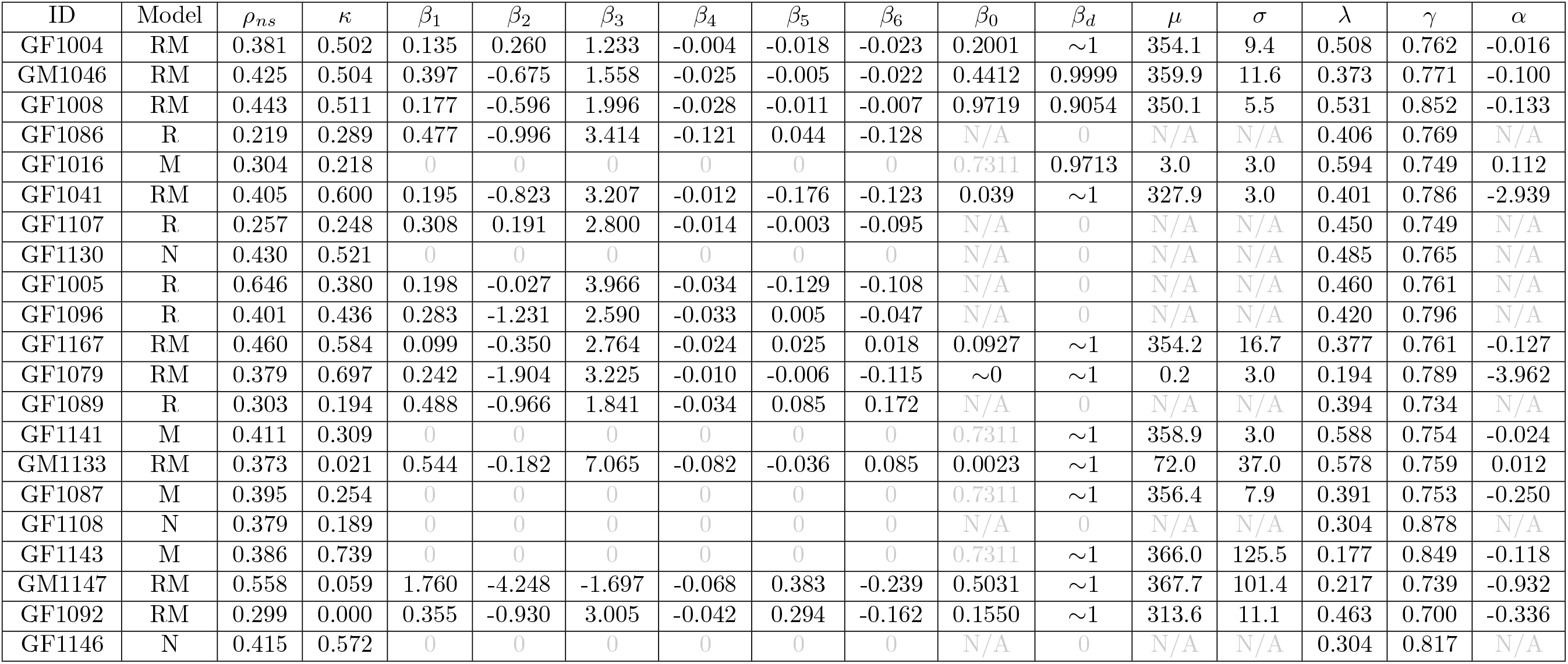
Parameter estimates for the “best model” (as identified by BIC) for each bear. Bears are listed in ascending order by number of GPS fixes. Note that the second letter of the bear ID indicates the sex of the individual. Gray text in the table indicates a parameter value that was fixed and not estimated for that model, and gray ”N/A” values indicate parameters that are not influential in the “best model” for that bear. Parameter estimates for *β*_0_ and *β_d_* that are very close to but not exactly 0 or 1 are indicated as such with a “ ”. See Tables **??** and **??** for 95% confidence intervals for each bear and each parameter.

Every bear spent more time in the non-stationary state than the stationary state, with estimates for λ being significantly lower than *γ*. The 95% confidence intervals for *γ*, the probability of remaining in the non-stationary state given the bear was already in it, were entirely above 0.5 for every member of the Mackenzie Delta population, implying that all bears were significantly more likely to stay in the non-stationary state than leave it at any given time. Conversely, 10 of the 21 bears had 95% confidence intervals for λ (concerning the stationary state) that were entirely below 0.5, and only one bear (GF1016) had a confidence interval for λ that was entirely above 0.5.

#### 4.2.2 Resource selection parameters

Of the 21 bears in the population, 14 (9 resource-memory + 5 resource-only) had resource selection in their “best model”. Some resource covariates displayed more within-population variation than others (Table 4). Only one (GF1167) of the 14 bears did not display significant selection for areas likely to contain berries (i.e., the 95% confidence interval for *β*_1_ was entirely above 0). The parameter estimate for GF1167 was positive but the lower confidence bound for *β*_1_ overlapped 0 (Table **??**). 8 of the 14 bears selected for areas closer to turbid water, suggesting attraction to riparian areas. None of the bears selected against this covariate. 11 of the 14 bears selected for areas indicative of high Arctic ground squirrel density, and only one bear (GM1147) displayed significant avoidance for these areas. Curiously, parameter estimates for *β*_4_, the selection coefficient for sweetvetch habitat, were negative for all 14 “resource-informed” bears. 10 of the 14 bears displayed significant selection against these areas. Bears generally displayed minimal responses to anthropogenic dwellings in the region. Only two (GF1005 and GF1041) bears displayed any significant pattern in relation to human settlements (both displayed selection), and only four displayed such behaviours with respect to industrial cabins (GF1089 avoided them while GF1005, GF1079, and GF1086 selected for areas closer to them).

#### 4.2.3 Memory parameters

13 of the 21 bears had memory incorporated in their “best model”, and most of these “memory-informed bears” returned to previously visited locations approximately 365 days after their last visit (Table 4). These trends were similar when resources were explicitly assumed to be seasonal (Table **??**), where the memory-only model provided the best explanation of most of the bears’ movements. Estimates of *β_d_* were often close to 1, and the 95% confidence intervals for this parameter never included 0.5, suggesting that memory played a part in movement for all of the memory-informed bears. 10 of the 13 bears had estimates for *μ* that were close to one year (>10 months or 300 days), implying that the majority of the population used a revisitation schedule of approximately one year. The other three bears (GF1016, GF1079, and GM1133) had estimates for *μ* of 3, 0.2, and 72 days, respectively (see the Discussion for more about GF1079). The median estimate value for *σ* was 9.4 days. For 9 of the 13 bears, the confidence interval for *σ* excluded 3 days (the lower optimization bound for *σ*; see Thompson et al. (2021) for more information), implying significant variation in the bears’ revisitation schedules. Estimates for *α* also varied between bears, ranging from −3.96 (GF1079) to 0.11 (GF1016). Based on the confidence intervals for this parameter, we found that 4 (GF1041, GF1079, GF1092, GM1147) of the 13 bears exhibited significantly heterogeneous perception of their landscapes, while 5 (GF1004, GF1016, GF1141, GF1167, GM1133) exhibited the opposite.

Of the 9 bears whose movements were best explained by the resource-memory model, 5 displayed especially selective revisitations to locations along their track (based on whether the 95% confidence interval for *β*_0_ was below 0.5). There was some individual variation in the estimates for *β*_0_ themselves, ranging from 0.00002 (GF1079) to 0.9719 (GF1008) (Table 4). However, the confidence interval for *β*_0_ was large for the latter. Figure 2 depicts an example of one of these 9 bears, GF1041, highlighted by a clear navigation to a previously visited location approximately a year later.

**Figure 2:**
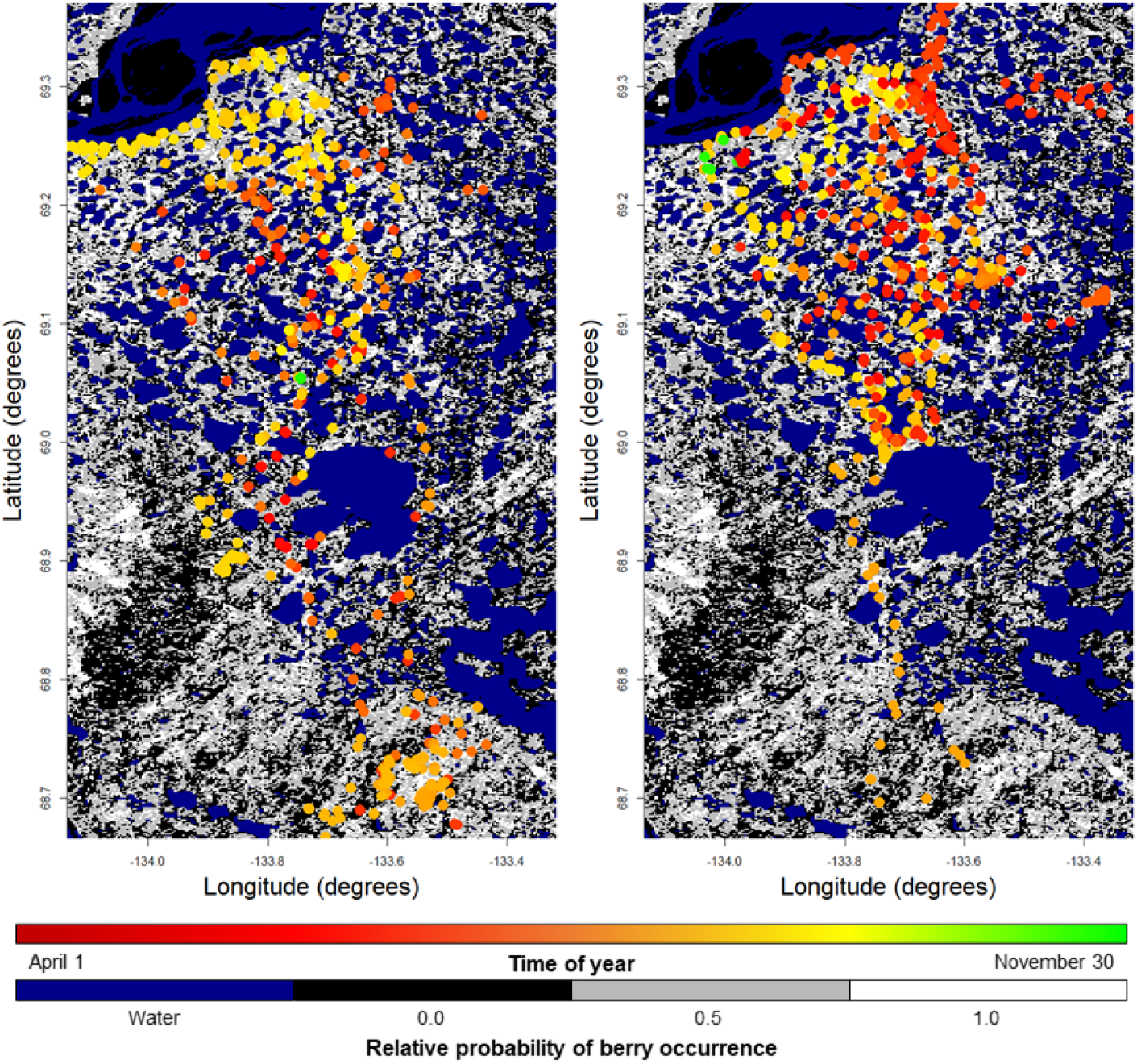
Movement track for bear ID GF1041 for the years 2003 (left) and 2004 (right). Each point on the animal’s track is colored according to the day of the year. The background displays values of the berry density layer (see the Appendix for details on how this layer was generated). Note the extended visitation of the southern part of the bear’s home range in 2003, followed by a directed navigation towards that same area at the same time in 2004. The movement patterns of GF1041 were best explained by the resource-memory model (Table 2).

## 5 Discussion

We used brown bear movement data from the Mackenzie Delta region to analyze the viability of a new model (Thompson et al., 2021) for an opportunistic omnivore living in a harsh and seasonal environment. This model incorporates a complex time-dependent spatial memory mechanism that allowed us to identify how long brown bears wait before returning to previously visited locations. We found a great deal of variation between individual bears, but this did not hinder our ability to observe population-level trends in the bears’ movement patterns. We also showed that representing resources as temporally constant was more effective in explaining the movement patterns of the brown bears than explicitly defining seasons for these resources.

The most common pattern observed in the population was a 365-day “circannual memory”, implying that many bears returned to portions of their home range that they visited roughly a year before (Figure 2). Previous work on this population identified a pattern of annual home range shift for Mackenzie Delta brown bears (Edwards et al., 2009). Potentially, brown bears maintain fidelity to portions of their home range, visiting those portions of the environment at the same time each year, and displaying less annual fidelity to other portions of their home range.

Some of the resource selection patterns observed in the “resource-informed” subset of the bear population could be explained by the nature of our landscape data. The lack of response to areas suggestive of sweetvetch presence was surprising, as almost every bear in the population displayed significant avoidance from such habitats. We based our estimate of sweetvetch presence on Porsild and Cody (1980), but other citations (e.g., Aiken et al. (2007)) indicate that they may appear closer to bodies of water, in sandy areas, or even in tundra (in fact, this could explain selection for areas closer to turbid water bodies such as rivers and coastlines).

Most bears did not display selection for or against anthropogenic structures, and when they did, the resulting behaviour was not always what we predicted. Bears are typically most affected by human presence when they have had a previous negative encounter with humans (Hertel et al., 2019), suggesting that this lack of significant selection could be explained by a lack of human-bear encounters. Many of the bears did not even go near Inuvik or Tuktoyaktuk while they were collared, suggesting a lack of encounters with humans.

When we adjusted our resource covariates such that they were explicitly assumed to only appear during a prescribed temporal interval, we found that the resource-only and resource-memory models were a significantly worse explanation of brown bear movement patterns. In fact, the number of bears that had resource selection included in their “best model” decreased from 14 with temporally constant resources to only 5 with explicitly seasonal resources. When resources were seasonally bounded, the memory-only model was much more effective than it was when resources were temporally constant. Recalling that the memory-only model is constructed independently of resources, this yields a clear ordering in the effectiveness of each model type for the entire population: resource-memory with constant resources > memory-only > resource-memory with seasonal resources. We do not dispute that these resources are indeed seasonal, but instead suggest that the landscape data we included in the model represent more than just the seasonal resources we included them for. For example, it may be possible that brown bears select for (and remember the location of) shrubby, berry-rich habitats outside of berry season, when they may provide other foraging benefits. Brown bears are opportunistic omnivores, and even when one food resource is widely available, they still maintain a balanced diet with multiple food sources (Robbins et al., 2007). When we restricted our resource availability to seasonal intervals, we nearly limited brown bear habitat selection to one resource covariate at a time, which may have been unrealistic. The correction we attempted in this model may not have worked for this population of bears, but in other systems (e.g., specialist herbivores in Kenyan savannas; Kartzinel and Pringle, 2020), it may be more appropriate.

The high inter-individual variation in the brown bear population is both interesting and unsurprising given what we know about the species and population. Brown bears undoubtedly possess the cognitive capability to remember the location of previously visited areas (Manchi and Swenson, 2005; Clapham et al., 2012; Selva et al., 2017; Wirsing et al., 2018), and the Arctic’s seasonal and spatial dynamics suggest that the spatio-temporal memory we tested here would be useful for optimal foraging (Fagan et al., 2013). It is then somewhat surprising that only 13 of the 21 bears in the population exhibited memory-informed movement according to our model selection process. One potential explanation is that if the temporal variation in the landscape is unpredictable, then periodic memory-informed movement may not improve foraging success (Mueller et al., 2011). Vegetation in the Mackenzie Delta region is somewhat unpredictable from year to year (Edwards et al., 2013), and while some resources may be available at the same place and time each year, memorizing the location of a patch that may not support resources in the future could be detrimental to foraging.

The model used here may not reliably be able to identify the correct signal with approximately one year of GPS collar data (Thompson et al., 2021), forcing us to question our results for bears with this much data. Although the simulation analyses performed with the model were effective for small data sizes, when Thompson et al. (2021) fit the model to individual years of data for the same bear, the “best model” often changed from year to year. This implies that either the bears are changing the way they move from one year to another, or that the model is unreliable in detecting a spatio-temporal memory signal without enough data. The former could arise as a result of reproductive activity in the population. When female bears have cubs, their movement strategies change as preventing infanticide and supporting their offspring become priorities (Edwards and Derocher, 2015). Male bears display much less behavioural plasticity with regard to reproduction, and all three of the male bears included in our analysis were best explained by the resource-memory model. Year-to-year variability in the landscape could also influence this behaviour; for example, if a bear finds food somewhere in one year, revisits that area 365 days later, and does not find food, it may use its cognitive map differently in subsequent years. Conversely, if a bear finds a new food source it may abandon its cognitive map and spend time at the newly found patch instead. We acknowledge that we cannot support or refute this hypothesis as an explanation for the within-individual variation we see between years, as it would require additional tracking data.

One bear, GF1079, yielded a set of unique parameter estimates from the resource-memory model as its “best model”. This bear had a *β*_0_ value close to 0 (approximately 0.000017) and its estimate for *μ* was less than 1 day, which implies that it was consistently moving away from locations it had visited very recently. This would be expected from an animal performing correlated random walk behaviour (its estimate for *κ* was 0.697, the second largest in the population), but in our model, we control for this behaviour by comparing observed steps to random steps simulated from a correlated random walk (as is done in traditional step selection analysis). This combination of parameter estimates only occurred with GF1079, which had only one year of location data (excluding the first year used for model training), suggesting that this occurrence is rare and may be alleviated with more data.

Our modelling framework focuses on behaviours that are observed in many other taxa, with potential for application in wildlife management. Boreal woodland caribou (*Rangifer tarandus caribou*) display site fidelity patterns that vary by season, displaying greater fidelity at different parts of the year (Lafontaine et al., 2017). Applying our framework to location data for woodland caribou, potentially breaking data up into seasonal partitions, would provide valuable inference about the extent of these patterns. The individual variation in brown bear movement behaviour was a key conclusion that we identified, which suggests that our modelling framework may be applicable to other ecological systems with high individual variation. As an example, black-legged kittiwakes (*Rissa tridactyla*) display individual differences in site fidelity when foraging near nesting colonies (Harris et al., 2020). These birds may not only exhibit different degrees of support for the resource-memory model, but the rate at which previous foraging paths are revisited (*μ*) may differ between individuals. Identifying the degree to which animals rely on memory is also important for translocation and reintroduction protocols. These protocols are often applied to animals that pose a high risk of coming into conflict with humans, transporting these animals to an environment they are unfamiliar with. Translocated animals that rely heavily on memory struggle to forage effectively in their new environments (Jesmer et al., 2018). Brown bears are frequently translocated, and these costly and time-consuming protocols significantly increase mortality risk if not executed properly (Milligan et al., 2018). These important and necessary decisions can be made more effectively with knowledge of how memory and familiarity impacts the movements of problem animals.

## 6 Acknowledgements

PRT was funded by the Ashley and Janet Cameron Graduate Scholarship, with support from UAlberta North, as well as the Alberta Graduate Excellence Scholarship. PRT was also supported by a University of Alberta Master’s Recruitment Award as well as a University of Alberta Doctoral Recruitment Award. MAL gratefully acknowledges support from a Canada Research Chair and NSERC Discovery grant. AED gratefully acknowledges support from an NSERC Discovery grant and from the Polar Continental Shelf Project of Natural Resources Canada. The authors declare no conflict of interest.

## 7 Author contributions

PRT designed the analysis with recommendations from MAL, AED, and MAE. MAE and AED collected and supplied brown bear data. PRT wrote the draft of the manuscript. All authors have reviewed and provided modifications to the manuscript.

