## Supplementary Information for "Time-dependent memory and individual variation in Arctic brown bears (*Ursus arctos*)"

### 1 Appendix: Supplementary Material

#### 1.1 Additional modelling details

We applied a discrete-time movement model developed by Thompson et al. (2021) that allows for inference on animal behaviour and cognition. We fit the model to data using maximum likelihood estimation, calculating the likelihood function as the product of individual likelihoods for each point (Thompson et al., 2021). The probability of taking a step is equal to the probability of taking that step from the non-stationary state ($f_{ns}$) multiplied by the probability of being in the non-stationary state, plus the the probability of taking that step from the stationary state ($f_s$) multiplied by the probability of being in the stationary state. Multiplying these probabilities together for each step yields the probability that the animal would produce a track exactly like the observed data under a given set of model parameters (Whoriskey et al., 2017). We optimized the negative logarithm of this function to maximize numerical precision, since these probabilities are always very, very small numbers.

We used automatic differentiation, via the Template Model Builder R package (Kristensen et al., 2016), to accurately fit the model amid high numerical complexity. For multidimensional functions such as ours, optimization routines typically require the calculation of the function's gradient. When the gradient of a function is 0 in every direction, there is potential that the optimizer has found a local optimum of that function. Traditional optimization does not require the user to provide a gradient along with their objective function, instead approximating the gradient at each point using finite-difference methods. Providing a gradient up front is faster computationally but difficult if there is no analytical expression for the gradient (as in most likelihood functions). TMB uses a pseudo-analytical automatic differentiation algorithm to calculate the gradient of a multidimensional likelihood function, alleviating the need for finite-difference approximation (Kristensen et al., 2016). While the negative log-likelihood function and its gradient are returned from TMB as R functions, the computational work is executed in C++, which also speeds up the optimization (Kristensen et al., 2016). We used the R *nlminb* function to optimize the appropriate TMB function for each model.

To prevent our model from producing errors or unrealistic results, we imposed bounds on some of the parameters. We bounded the estimation for $\mu$ at 365 days, the amount of GPS data we set aside for "training" the bears' memory. If $\mu$ were to be greater than this value, there would not be a testable memory signal for some of the data (i.e., we would not always know where the bear was $\mu$ days before). To identify potential signals greater than 365 days, we would have to sacrifice bears with insufficient data, thus decreasing our "population size" and our statistical power. We performed the optimization routine twice for each bear with different "initial values" for $\mu$, since there was potential for local optima along the axis of this parameter (Thompson et al., 2021). We also applied a lower optimization bound of 3 days (18 GPS fixes) on $\sigma$. When $\sigma < 3$ days, the partial derivative of our likelihood function with respect to $\mu$

became noisy, leading to computational errors in optimization. We additionally required $\alpha < -\log_{10} \frac{30}{1000} \approx 1.52$. We chose this bound because each grid cell is $\frac{30}{1000}$ km in length. For parameters with fairly restrictive bounds ($\lambda$, $\gamma$, $\beta_d$, and $\beta_0$, which are bounded between 0 and 1), we performed logit transformations ($\tilde{\lambda} = \log \frac{1}{1-\lambda}$, for example) so the optimizer would more effectively traverse the parameter space.

Instead of using traditional Wald-type confidence intervals for our parameters, we found them explicitly using likelihood profiling techniques (Fischer and Lewis, 2020). The algorithm used here finds confidence intervals for one parameter at a time by performing a binary search algorithm for a target function value (the optimal function value minus a confidence threshold of 1.96). The algorithm searches for each parameter by moving away from the optimal value in parameter space, fixing that parameter and optimizing the other ones. This process goes on until the algorithm gets close enough to the target function value. This means that, unlike traditional confidence intervals, the lower confidence bound may be farther away from the parameter estimate than the upper confidence bound.

#### 1.2 Generating the berry density raster

We generated a vegetation class raster using the decision tree from Ducks Unlimited (2002) to classify each 30 x 30 m grid cell in the Mackenzie Delta region into one of 46 classes. Table S1 describes each class as well as its value for the berry probability raster used in the model.

#### 1.3 Supplementary tables

| Description | Berry probability | Percent of landscape |
|---|---|---|
| Open needleleaf | 0.5 | 0.397 |
| Woodland needleleaf (other) | 1 | 1.481 |
| Woodland needleleaf (moss) | 1 | 0.303 |
| Woodland needleleaf (lichen) | 1 | 0.524 |
| Closed deciduous | 0.5 | 0.082 |
| Open deciduous | 0.5 | 0.344 |
| Closed tall shrub | 0 | 1.476 |
| Open tall shrub | 0 | 0.007 |
| Medium willow shrub | 0 | 0.480 |
| Medium-tall willow shrub | 0 | 0.356 |
| Medium-tall shrub (other) | 0 | 0.027 |
| Low shrub (other) | 0 | 14.087 |
| Low shrub (recently burned) | 0 | 0.352 |
| Low shrub (wet) | 0 | 0.004 |
| Low shrub (floodplain) | 0 | 0.688 |
| Low shrub (delta lowlands) | 0 | 1.243 |
| Low shrub - wet graminoid | 0 | 0.012 |
| Low shrub - tussock | 0 | 1.135 |
| Low shrub (upland) | 0.5 | 0.368 |
| Dwarf shrub (other) | 1 | 9.097 |
| Dwarf shrub (tussock) | 0 | 5.331 |
| Dwarf shrub (lichen) | 1 | 2.239 |
| Dwarf shrub (unknown - Kendall area) | 0.5 | 0.046 |
| Dwarf shrub (tussock/Dryas) | 0 | 0.072 |
| Dwarf shrub (Dryas/heather) | 0.5 | 0.796 |
| Dwarf shrub (sloped hummocks) | 0 | 0.035 |
| Dwarf shrub (wet graminoid) | 0.5 | 0.160 |
| Tussock tundra | 0.5 | 1.407 |
| Lichen | 0.5 | 0.243 |
| Wet graminoid (wetland depressions or lake edges) | 0 | 1.762 |
| Wet graminoid (northern delta) | 0 | 1.429 |
| Wet graminoid (floodplain) | 0 | 1.101 |
| Wet graminoid (some shrubs) | 0 | 0.111 |
| Wet graminoid (unknown) | 0 | 0.003 |
| Dwarf shrub mosaic (wet) | 0.5 | 0.355 |
| Dwarf shrub mosaic (very wet) | 0.5 | 0.761 |
| Aquatic bed | 0 | 0.014 |
| Emergent vegetation | 0 | 1.002 |
| Emergent and other wet wetland areas | 0 | 1.424 |
| Clear water | 0 | 20.108 |
| Turbid water | 0 | 25.385 |
| Sparse vegetation | 0 | 0.394 |
| Sparse / non-vegetated (unsure) | 0 | 0.030 |
| Non-vegetated | 0 | 0.670 |
| Other | 0 | 0.001 |
| Unclassified | 0 | 2.655 |



Table S1: Classification method used to generate the berry density raster for the resource-only and resource-memory models. The name of each original vegetation class is shown along with the assigned probability, representing how correlated each habitat type is to the presence of berries, as well as the percentage of the landscape that is covered by each class.

| ID | Model | $\rho_{ns}$ | $\kappa$ | $\beta_1$ | $\beta_2$ | $\beta_3$ | $\beta_4$ | $\beta_5$ | $\beta_6$ | $\beta_0$ | $\beta_d$ | $\mu$ | $\sigma$ | $\lambda$ | $\gamma$ | $\alpha$ |
|---|---|---|---|---|---|---|---|---|---|---|---|---|---|---|---|---|
| GF1004 | RM | 0.356 | 0.404 | 0.065 | -0.008 | 0.681 | -0.015 | -0.055 | -0.101 | 0.1262 | ~1 | 351.0 | 7.2 | 0.459 | 0.731 | -0.052 |
| GM1046 | RM | 0.397 | 0.404 | 0.300 | -1.212 | 0.989 | -0.041 | -0.066 | -0.041 | 0.2195 | 0.5004 | 355.7 | 7.9 | 0.315 | 0.740 | -0.341 |
| GF1008 | RM | 0.414 | 0.410 | 0.099 | -1.052 | 1.473 | -0.044 | -0.085 | -0.085 | 0.2010 | 0.5000 | 345.0 | 3.3 | 0.467 | 0.825 | -0.359 |
| GF1086 | R | 0.205 | 0.191 | 0.353 | -1.428 | 2.195 | -0.226 | -0.069 | -0.241 | N/A | 0 | N/A | N/A | 0.348 | 0.737 | N/A |
| GF1016 | M | 0.281 | 0.106 | 0 | 0 | 0 | 0 | 0 | 0 | 0.7311 | 0.9168 | 1.3 | 3.0 | 0.542 | 0.712 | 0.067 |
| GF1041 | RM | 0.376 | 0.488 | 0.122 | -1.393 | 2.517 | -0.024 | -0.333 | -0.281 | 0.0127 | ~1 | 326.9 | 3.0 | 0.340 | 0.753 | -5.056 |
| GF1107 | R | 0.237 | 0.130 | 0.192 | -0.569 | 2.385 | -0.027 | -0.220 | -0.314 | N/A | 0 | N/A | N/A | 0.385 | 0.710 | N/A |
| GF1130 | N | 0.395 | 0.393 | 0 | 0 | 0 | 0 | 0 | 0 | N/A | 0 | N/A | N/A | 0.419 | 0.726 | N/A |
| GF1005 | R | 0.591 | 0.25 | 0.111 | -0.691 | 2.676 | -0.062 | -0.212 | -0.192 | N/A | 0 | N/A | N/A | 0.391 | 0.720 | N/A |
| GF1096 | R | 0.367 | 0.306 | 0.183 | -1.840 | 1.755 | -0.057 | -0.088 | -0.144 | N/A | 0 | N/A | N/A | 0.344 | 0.758 | N/A |
| GF1167 | RM | 0.417 | 0.437 | -0.003 | -0.730 | 1.765 | -0.043 | -0.063 | -0.073 | 0.0392 | ~1 | 346.8 | 10.2 | 0.300 | 0.716 | -0.184 |
| GF1079 | RM | 0.343 | 0.541 | 0.141 | -2.581 | 2.450 | -0.028 | -0.091 | -0.211 | ~0 | 0.5022 | 0.2 | 3.0 | 0.117 | 0.744 | -8.007 |
| GF1089 | R | 0.272 | 0.041 | 0.306 | -1.683 | -0.432 | -0.070 | -0.053 | 0.023 | N/A | 0 | N/A | N/A | 0.309 | 0.682 | N/A |
| GF1141 | M | 0.356 | 0.099 | 0 | 0 | 0 | 0 | 0 | 0 | 0.7311 | ~1 | 357.5 | 3.0 | 0.488 | 0.685 | -0.0680 |
| GM1133 | RM | 0.323 | 0.000 | 0.129 | -1.158 | -0.309 | -0.659 | -0.257 | -0.140 | ~0 | ~1 | 54.5 | 28.1 | 0.476 | 0.689 | -0.137 |
| GF1087 | M | 0.340 | 0.038 | 0 | 0 | 0 | 0 | 0 | 0 | 0.7311 | ~1 | 350.9 | 26.8 | 0.277 | 0.680 | -0.548 |
| GF1108 | N | 0.328 | 0.000 | 0 | 0 | 0 | 0 | 0 | 0 | N/A | 0 | N/A | N/A | 0.142 | 0.820 | N/A |
| GF1143 | M | 0.330 | 0.493 | 0 | 0 | 0 | 0 | 0 | 0 | 0.7311 | ~1 | 199.1 | 3.0 | 0.059 | 0.781 | -0.374 |
| GM1147 | RM | 0.465 | 0.000 | 1.304 | -5.949 | -2.186 | -0.188 | -0.537 | -0.426 | 0.2917 | ~1 | 235.8 | 72.5 | 0.101 | 0.649 | -1.136 |
| GF1092 | RM | 0.243 | 0.000 | 0.035 | -3.423 | 1.992 | -0.084 | -0.141 | -0.560 | 0.0534 | ~1 | 308.1 | 6.9 | 0.315 | 0.592 | -0.529 |
| GF1146 | N | 0.321 | 0.181 | 0 | 0 | 0 | 0 | 0 | 0 | N/A | 0 | N/A | N/A | 0.110 | 0.698 | N/A |

Table S2a: 95% lower confidence bounds for the "best model" (as identified by BIC) for each bear, calculated using likelihood profiling. Bears are listed in ascending order by number of GPS fixes. Gray text in the table indicates a parameter value that was fixed and not estimated for that model, and gray "N/A" values indicate parameters that are not influential in the "best model" for that bear. Parameter estimates for $\beta_0$ and $\beta_d$ that are very close to but not exactly 0 or 1 are indicated as such with a " ". See Table **??** for descriptions of each parameter.

| ID | Model | $\rho_{ns}$ | $\kappa$ | $\beta_1$ | $\beta_2$ | $\beta_3$ | $\beta_4$ | $\beta_5$ | $\beta_6$ | $\beta_0$ | $\beta_d$ | $\mu$ | $\sigma$ | $\lambda$ | $\gamma$ | $\alpha$ |
|---|---|---|---|---|---|---|---|---|---|---|---|---|---|---|---|---|
| GF1004 | RM | 0.407 | 0.603 | 0.204 | 0.525 | 1.763 | 0.005 | 0.091 | 0.054 | 0.3049 | ~1 | 357.1 | 11.9 | 0.557 | 0.791 | -0.016 |
| GM1046 | RM | 0.455 | 0.607 | 0.494 | -0.154 | 2.104 | -0.010 | 0.057 | 0.085 | ~1 | ~1 | 365.0 | 18.0 | 0.433 | 0.800 | -0.100 |
| GF1008 | RM | 0.475 | 0.613 | 0.254 | -0.151 | 2.493 | -0.013 | 0.062 | 0.084 | ~1 | ~1 | 353.1 | 10.0 | 0.594 | 0.877 | -0.133 |
| GF1086 | R | 0.235 | 0.390 | 0.599 | -0.573 | 4.564 | -0.044 | 0.157 | -0.014 | N/A | 0 | N/A | N/A | 0.465 | 0.799 | N/A |
| GF1016 | M | 0.329 | 0.330 | 0 | 0 | 0 | 0 | 0 | 0 | 0.7311 | 0.9910 | 4.6 | 4.1 | 0.644 | 0.783 | 0.112 |
| GF1041 | RM | 0.437 | 0.714 | 0.266 | -0.268 | 3.882 | -0.001 | -0.018 | 0.036 | 0.0779 | ~1 | 328.9 | 3.4 | 0.464 | 0.816 | -1.869 |
| GF1107 | R | 0.280 | 0.368 | 0.425 | 0.932 | 3.215 | -0.003 | 0.211 | 0.123 | N/A | 0 | N/A | N/A | 0.516 | 0.786 | N/A |
| GF1130 | N | 0.469 | 0.651 | 0 | 0 | 0 | 0 | 0 | 0 | N/A | 0 | N/A | N/A | 0.551 | 0.801 | N/A |
| GF1005 | R | 0.707 | 0.511 | 0.285 | 0.620 | 5.169 | -0.010 | -0.043 | -0.022 | N/A | 0 | N/A | N/A | 0.531 | 0.799 | N/A |
| GF1096 | R | 0.439 | 0.566 | 0.382 | -0.645 | 3.384 | -0.013 | 0.096 | 0.048 | N/A | 0 | N/A | N/A | 0.498 | 0.832 | N/A |
| GF1167 | RM | 0.508 | 0.735 | 0.202 | 0.019 | 3.714 | -0.007 | 0.114 | 0.109 | 0.1951 | ~1 | 362.7 | 25.6 | 0.458 | 0.802 | -0.127 |
| GF1079 | RM | 0.420 | 0.856 | 0.343 | -1.265 | 3.982 | 0.007 | 0.102 | -0.020 | 0.0017 | ~1 | 0.6 | 3.3 | 0.286 | 0.830 | -1.783 |
| GF1089 | R | 0.338 | 0.349 | 0.673 | -0.279 | 3.861 | -0.004 | 0.222 | 0.321 | N/A | 0 | N/A | N/A | 0.481 | 0.782 | N/A |
| GF1141 | M | 0.477 | 0.523 | 0 | 0 | 0 | 0 | 0 | 0 | 0.7311 | ~1 | 360.3 | 3.9 | 0.684 | 0.816 | -0.024 |
| GM1133 | RM | 0.433 | 0.229 | 0.967 | 0.759 | 14.020 | 0.124 | 0.190 | 0.305 | 0.4537 | ~1 | 79.3 | 48.1 | 0.676 | 0.821 | 0.014 |
| GF1087 | M | 0.461 | 0.473 | 0 | 0 | 0 | 0 | 0 | 0 | 0.7311 | ~1 | 361.4 | 15.9 | 0.513 | 0.818 | -0.053 |
| GF1108 | N | 0.440 | 0.399 | 0 | 0 | 0 | 0 | 0 | 0 | N/A | 0 | N/A | N/A | 0.494 | 0.925 | N/A |
| GF1143 | M | 0.455 | 0.995 | 0 | 0 | 0 | 0 | 0 | 0 | 0.7311 | ~1 | 366.0 | 349.7 | 0.353 | 0.904 | -0.118 |
| GM1147 | RM | 0.676 | 0.326 | 2.259 | -2.749 | -1.251 | 0.006 | 0.54 | 0.097 | 0.5068 | ~1 | 367.7 | 139.8 | 0.371 | 0.818 | -0.6730 |
| GF1092 | RM | 0.373 | 0.110 | 0.666 | 1.387 | 4.021 | -0.006 | 0.726 | 0.250 | 0.7664 | ~1 | 322.7 | 19.3 | 0.613 | 0.796 | -0.114 |
| GF1146 | N | 0.549 | 0.985 | 0 | 0 | 0 | 0 | 0 | 0 | N/A | 0 | N/A | N/A | 0.556 | 0.906 | N/A |

Table S2b: 95% upper confidence bounds for the “best model” (as identified by BIC) for each bear, calculated using likelihood profiling. Bears are listed in ascending order by number of GPS fixes. Gray text in the table indicates a parameter value that was fixed and not estimated for that model, and gray ”N/A” values indicate parameters that are not influential in the “best model” for that bear. Parameter estimates for $\beta_0$ and $\beta_d$ that are very close to but not exactly 0 or 1 are indicated as such with a “ ”. See Table **??** for descriptions of each parameter.

| ID | Model | $\rho_{ns}$ | $\kappa$ | $\beta_1$ | $\beta_2$ | $\beta_3$ | $\beta_4$ | $\beta_5$ | $\beta_6$ | $\beta_0$ | $\beta_d$ | $\mu$ | $\sigma$ | $\lambda$ | $\gamma$ | $\alpha$ |
|---|---|---|---|---|---|---|---|---|---|---|---|---|---|---|---|---|
| GF1004 | RM | 0.383 | 0.502 | 0.265 | 0.300 | -0.265 | -0.015 | 0.068 | 0.007 | 0.4360 | ~1 | 0.8 | 3.0 | 0.508 | 0.759 | -1.611 |
| GM1046 | M | 0.428 | 0.517 | 0 | 0 | 0 | 0 | 0 | 0 | 0.7311 | ~1 | 359.1 | 10.2 | 0.390 | 0.772 | -0.126 |
| GF1008 | M | 0.443 | 0.526 | 0 | 0 | 0 | 0 | 0 | 0 | 0.7311 | ~1 | 351.6 | 4.4 | 0.528 | 0.852 | -0.133 |
| GF1086 | R | 0.219 | 0.229 | 0.945 | -0.792 | 3.529 | -0.130 | 0.079 | -0.120 | N/A | 0 | N/A | N/A | 0.403 | 0.770 | N/A |
| GF1016 | M | 0.301 | 0.220 | 0 | 0 | 0 | 0 | 0 | 0 | 0.7311 | 0.9782 | 3.0 | 3.0 | 0.601 | 0.758 | 0.112 |
| GF1041 | N | 0.405 | 0.578 | 0 | 0 | 0 | 0 | 0 | 0 | N/A | 0 | N/A | N/A | 0.399 | 0.785 | N/A |
| GF1107 | M | 0.258 | 0.253 | 0 | 0 | 0 | 0 | 0 | 0 | 0.7311 | ~1 | 365.0 | 65.0 | 0.448 | 0.750 | 0.140 |
| GF1130 | RM | 0.430 | 0.529 | 0.178 | -0.110 | -0.325 | -0.051 | 0.002 | -0.031 | 0.4415 | ~1 | 0.8 | 3.0 | 0.488 | 0.768 | -1.762 |
| GF1005 | M | 0.651 | 0.331 | 0 | 0 | 0 | 0 | 0 | 0 | 0.7311 | 0.9953 | 5.6 | 3.0 | 0.473 | 0.758 | -0.256 |
| GF1096 | M | 0.400 | 0.402 | 0 | 0 | 0 | 0 | 0 | 0 | 0.7311 | ~1 | 348.1 | 26.9 | 0.405 | 0.794 | -0.079 |
| GF1167 | M | 0.463 | 0.574 | 0 | 0 | 0 | 0 | 0 | 0 | 0.7311 | ~1 | 364.8 | 6.3 | 0.387 | 0.761 | -0.126 |
| GF1079 | R | 0.378 | 0.728 | 0.376 | -2.161 | 2.508 | 0.012 | -0.059 | -0.141 | N/A | 0 | N/A | N/A | 0.171 | 0.786 | N/A |
| GF1089 | R | 0.298 | 0.192 | 1.506 | -0.783 | 1.243 | -0.005 | 0.056 | 0.196 | N/A | 0 | N/A | N/A | 0.379 | 0.740 | N/A |
| GF1141 | M | 0.419 | 0.529 | 0 | 0 | 0 | 0 | 0 | 0 | 0.7311 | ~1 | 365.0 | 11.9 | 0.606 | 0.754 | -0.023 |
| GM1133 | M | 0.374 | 0.080 | 0 | 0 | 0 | 0 | 0 | 0 | 0.7311 | ~1 | 54.5 | 26.7 | 0.595 | 0.768 | -0.212 |
| GF1087 | M | 0.400 | 0.295 | 0 | 0 | 0 | 0 | 0 | 0 | 0.7311 | ~1 | 357.9 | 9.2 | 0.396 | 0.744 | -0.338 |
| GF1108 | N | 0.376 | 0.211 | 0 | 0 | 0 | 0 | 0 | 0 | N/A | 0 | N/A | N/A | 0.273 | 0.880 | N/A |
| GF1143 | M | 0.388 | 0.722 | 0 | 0 | 0 | 0 | 0 | 0 | 0.7311 | ~1 | 352.8 | 15.1 | 0.179 | 0.848 | -0.119 |
| GM1147 | M | 0.556 | 0.115 | 0 | 0 | 0 | 0 | 0 | 0 | 0.7311 | ~1 | 260.9 | 3.0 | 0.204 | 0.742 | -0.477 |
| GF1092 | M | 0.301 | 0.000 | 0 | 0 | 0 | 0 | 0 | 0 | 0.7311 | ~1 | 363.1 | 42.0 | 0.469 | 0.714 | 0.106 |
| GF1146 | M | 0.412 | 0.549 | 0 | 0 | 0 | 0 | 0 | 0 | 0.7311 | 0.9998 | 350.0 | 3.0 | 0.286 | 0.819 | -0.162 |

Table S3: Parameter estimates for the "best model" (as identified by BIC) for each bear with explicit expression of resource seasonality. Bears are listed in ascending order by number of GPS fixes. Note that the second letter of the bear ID indicates the sex of the individual. Gray text in the table indicates a parameter value that was fixed and not estimated for that model, and gray "N/A" values indicate parameters that are not influential in the "best model" for that bear. Parameter estimates for $\beta_0$ and $\beta_d$ that are very close to but not exactly 0 or 1 are indicated as such with a " ".
